# A Machine Learning Framework for Serogroup Classification of pathogenic species of *Leptospira* Based on *rfb* Locus Profiles

**DOI:** 10.64898/2026.03.04.708288

**Authors:** Edson de Carvalho Ferreira Filho, Pedro Melo Arruda, Leonardo Cabral Afonso Ferreira, Maria Raquel Venturim Cosate, Tetsu Sakamoto

**Affiliations:** Bioinformatics Multidisciplinary Environment (BioME), Instituto Metrópole Digital (IMD), Universidade Federal do Rio Grande do Norte (UFRN), Brazil; Independent Researcher, Boston, Massachusetts, USA

**Keywords:** Leptospirosis, Serotyping, O-antigen, Lipopolysaccharide, Epidemiology, Random Forest

## Abstract

*Leptospira* is a highly diverse genus traditionally classified by serological assays into more than 30 serogroups and over 300 serovars. However, this classification system is often complex and inconsistent, as cross-reactions between antigens can lead to ambiguous results. Moreover, serological tests such as MAT and CAAT are labor-intensive, require live cultures, and are difficult to standardize across laboratories. To overcome these limitations, we compiled genomic data from 721 pathogenic *Leptospira* samples obtained from NCBI RefSeq and BIGSdb (Institut Pasteur) to develop a machine learning framework capable of predicting serological classification directly from genomic information. Our approach focuses on the *rfb* locus, a genomic region associated with lipopolysaccharide biosynthesis and antigenic diversity, and was designed to operate in two stages: the first stage assigns samples to one of four major serological classes, while the second stage classifies them into their respective serogroups. Models from both classification stages achieved high predictive performance, with perfect score in the first and a mean F1-score of 0.948 in the second stage. Feature importance analysis revealed non-random clustering of highly informative genes within the *rfb* locus and demonstrated that serogroup discrimination is driven by combinatorial patterns of gene presence and absence. Based on the strong genetic coherence observed at the *rfb* locus and shared antigenic features, we propose the term “seroclass” to designate these higher-order groupings. This approach provides a scalable and reproducible alternative to traditional serological testing and offers valuable applications for epidemiological surveillance, outbreak investigation, and vaccine development within the *Leptospira* genus.

**Highlights:**

- Genomic data enable accurate inference of *Leptospira* serological classification.
- The approach predicts serogroups directly from *rfb* locus gene composition.
- The model provides a scalable alternative to labor-intensive serological assays.
- We introduce the term “seroclass”, a higher level of serological organization.
- The proposed framework supports epidemiological surveillance and vaccine design.

## 1. Introduction

The *Leptospira* genus comprises spirochete bacteria (Phylum Spirochaetes) characterized by their helical morphology and high motility (Ko et al., 2009; Levett, 2001). Pathogenic species of *Leptospira* are responsible for leptospirosis, one of the most widespread zoonotic diseases worldwide (Costa et al., 2015). Infection occurs through contact with water or soil contaminated with the urine of infected animals, leading to a wide spectrum of clinical outcomes ranging from mild febrile illness to severe Weil’s disease and pulmonary hemorrhage (Haake and Levett, 2015).

There are two main approaches for classifying *Leptospira* isolates. The first is taxonomic classification, which is based on the evolutionary relationships among isolates. The advent of molecular and genomic methods prompted a taxonomic reorganization of the genus *Leptospira*. Initially divided into “pathogenic” and “saprophytic” species (Levett, 2001), the genus is now recognized to comprise at least 77 species (Dos Santos Ribeiro et al., 2023; Fernandes et al., 2022; Hamond et al., 2025; Korba et al., 2021), distributed across four phylogenetic clades with distinct ecological and pathogenic characteristics: P1 (pathogenic *sensu stricto*), P2 (intermediate), S1 (saprophytic *sensu stricto*), and S2 (environmental) (Vincent et al., 2019). Pathogenic species such as *L. interrogans, L. kirschneri*, and *L. borgpetersenii* are primarily associated with human and animal disease, whereas intermediate and saprophytic species are mainly found in environmental niches.

The second approach is serological classification, which relies on antigenic differences among isolates. *Leptospira* classification has traditionally been based on serological tests, primarily the microscopic agglutination test (MAT) and the cross-agglutination absorption test (CAAT), which led to the description of more than 300 serovars grouped into 30 serogroups (Caimi and Ruybal, 2020; Faine et al., 1999; Picardeau, 2017). It is important to note that serological classification does not necessarily reflect genetic relatedness, as distinct species may contain strains belonging to the same serogroup, and conversely, a single species may comprise multiple serogroups (Faine et al., 1999; Kalambaheti et al., 1999; Mitchison et al., 1997). Accurate identification of *Leptospira* strains and their serogroups or serovars is crucial for understanding epidemiological dynamics, tracking outbreaks, and supporting vaccine development (Levett, 2001; Picardeau, 2017). Although MAT and CAAT are considered the gold standard (Terpstra et al., 2003), they are laborious, require maintaining large reference collections, and are prone to cross-reactivity and subjective interpretation.

The development of alternative methods for serological classification is therefore of great interest in the field. In this context, molecular approaches have emerged as promising tools to complement traditional serological assays (Ciurariu et al., 2025; Giraud-Gatineau et al., 2025). At the molecular level, the *rfb* locus, which encodes enzymes involved in the biosynthesis of the O-antigen component of lipopolysaccharides (LPS), is largely responsible for *Leptospira*’s antigenic diversity and therefore underlies serogroup and serovar classification (de la Peña-Moctezuma et al., 2001, 1999). Variation in gene content and organization within the *rfb* cluster determines structural differences in LPS, leading to distinct serological profiles among strains. Consequently, the *rfb* region represents a genetic link between traditional serological classification and genomic data.

Despite significant advances in sequencing and comparative genomics (Chinchilla et al., 2023; Ferreira et al., 2024; Medeiros et al., 2022; Nieves et al., 2022), connecting the extensive genomic diversity of *Leptospira* to its antigenic properties remains challenging. Machine learning (ML) methods have emerged as powerful tools for uncovering complex, nonlinear patterns in high-dimensional genomic data, enabling phenotype prediction such as antimicrobial resistance, host specificity, or serotype (Drouin et al., 2019; Lees et al., 2020). These approaches can detect subtle genomic signatures that are not easily captured by phylogenetic trees or pairwise similarity analyses. However, ML-based serogroup prediction in *Leptospira* remains largely unexplored.

Here, we propose a machine learning framework to classify *Leptospira* isolates into serogroups using genome-derived features. By integrating supervised learning algorithms with genomic representations, we aim to evaluate whether genomic variation—particularly in loci such as *rfb*—can reliably predict serogroup identity. Our results demonstrate that genomic data can reproduce and extend traditional serological classifications, providing a scalable and objective approach for *Leptospira* characterization and surveillance.

## 2. Materials and Methods

### 2.1 Genome and serological dataset collection

*Leptospira* genome sequences were obtained from the NCBI RefSeq Genomes database (Goldfarb et al., 2025) and the BIGSdb–Institut Pasteur repository (Jolley and Maiden, 2010) in October, 2024. Serological annotations were also retrieved from the corresponding databases. Only non-mutant isolates of the P1 group, with genome assemblies comprising fewer than 300 contigs and with available serogroup annotations, were included in the analysis. Potential redundancies between the two databases were identified and removed. Consistency of serogroup annotation between NCBI and BIGSdb entries was verified, and any discrepancies were resolved through manual curation and cross-checking with the literature. Additionally, we removed the sample *L. interrogans* str. UT053 (srg. Sejroe, srv. Medanensis) from the dataset, as it has been previously reported to deviate from the canonical *rfb* locus organization (Ferreira et al., 2024). Following these procedures, 721 *Leptospira* genomes were retained for downstream analyses (Supplementary Table S1). Additionally, we retrieved 30 genomic samples from NCBI RefSeq Genomes with serological information that were published after the development of the classification models to serve as validation dataset (Supplementary Table S2).

### 2.2 Feature matrix construction based on the *rfb* locus

The feature matrix used as input for the machine learning model comprised the identity values shared by each genome sample with the genes of the *rfb* locus as features (Supplementary Table S3). To construct the matrix, we first selected one representative strain from each *Leptospira* serogroup (Table 1), prioritizing genomes of high sequencing quality and that contained the full *rfb* locus assembled within a single contig. From each selected genome, amino acid sequences corresponding to the *rfb* locus genes were extracted. The resulting set of all protein sequences extracted was then clustered using CD-HIT (Fu et al., 2012) with a similarity threshold of 80% to reduce redundancy. The clustering analysis resulted in a total of 592 clusters. Representative sequences from each cluster with at least 100 amino acids long, totaling 549 proteins, were retrieved and used as queries in TBLASTN (Altschul et al., 1990) searches against the genome assemblies of the isolates in the dataset. For each query–subject comparison, the percentage of amino acid identity of the best hit was corrected by the query length to account for partial alignments. The corrected identity values were used to fill the numerical feature matrix, which served as input for training the machine learning classification models.

**Table 1:** Representative leptospiral samples of each serogroup.

| Seroclass | Serogroup | Genome accession | Sample |
| --- | --- | --- | --- |
| I | Ballum | GCF_009884235.1 | *Leptospira borgpetersenii* str. R14L |
|  | Canicola | GCF_008831465.1 | *Leptospira interrogans* str. 611 |
|  | Celledoni | GCF_022344045.1 | *Leptospira weilii* str. FMAS_PD2 |
|  | Icterohaemorrhagiae | GCF_000231175.1 | *Leptospira interrogans* serovar Lai str. IPAV |
|  | Javanica | GCF_024722275.1 | *Leptospira borgpetersenii* str. I2 |
|  | Manhao | GCF_000243815.2 | *Leptospira alexanderi* serovar Manhao 3 str. L 60 |
|  | Pyrogenes | GCF_022559725.1 | *Leptospira interrogans* str. FMAS_KG2 |
|  | Ranarum | GCF_000332415.1 | *Leptospira weilii* serovar Ranarum str. ICFT |
|  | Sarmin | GCF_030023765.1 | *Leptospira santarosai* str. MMD3 |
| II | Australis | GCF_022819715.1 | *Leptospira noguchii* str. IP1611024 |
|  | Autumnalis | GCF_022819425.1 | *Leptospira noguchii* str. IP1709037 |
|  | Cynopteri | GCF_000243695.2 | *Leptospira kirschneri* serovar Cynopteri str. 3522 CT |
|  | Djasiman | GCF_000216195.1 | *Leptospira interrogans* serovar Djasiman str. LT1649 |
|  | Grippotyphosa | GCF_024526015.1 | *Leptospira kirschneri* str. FMAS_PN5 |
|  | Panama | GCF_000306255.2 | *Leptospira noguchii* serovar Panama str. CZ214 |
|  | Pomona | GCF_001857845.1 | *Leptospira interrogans* serovar Pomona str. AKRFB |
| III | Hebdomadis | GCF_000244115.1 | *Leptospira interrogans* serovar Hebdomadis str. R499 |
|  | Mini | GCF_000306675.2 | *Leptospira mayottensis* str. 200901116 |
|  | Sejroe | GCF_003254845.1 | *Leptospira borgpetersenii* serovar Hardjo-bovis str. 203 |
| IV | Bataviae | GCF_014858865.1 | *Leptospira interrogans* serovar Bataviae str. 1489 |
|  | Shermani | GCF_000332395.2 | *Leptospira santarosai* serovar Shermani str. 1342KT |
|  | Tarassovi | GCF_024704545.1 | *Leptospira borgpetersenii* str. MN900 |

To further minimize redundancy among samples, we performed a clustering analysis based on Pearson correlation coefficients computed from the feature matrix (Supplementary Table S4). Samples exhibiting extremely high correlation (ρ > 0.999) were grouped into clusters, and a single representative genome from each cluster was retained for subsequent analyses (Supplementary Table S1). This stringent threshold was adopted because isolates belonging to the same serogroup, even when originating from different geographic locations or years of isolation, frequently exhibited correlation values approaching 0.99. Nearly all clusters generated through this procedure contained isolates from the same species and serogroup. The only exception was a cluster comprising three isolates from serogroup Icterohaemorrhagiae and two from serogroup Javanica; these samples were excluded from further analyses to avoid potential misclassification bias. After this final filtering step, the dataset consisted of 384 *Leptospira* genomes (rows) and 549 *rfb*-encoded protein features (columns).

### 2.3 Exploratory Data Analysis and Visualization

To explore the structure and variability of the genomic feature matrix, a principal component analysis (PCA) was performed using the *scikit-learn* package (Pedregosa et al., 2011). PCA was applied to reduce the dimensionality of the dataset and to visualize the distribution of *Leptospira* isolates in a two-dimensional space. Data visualization was carried out in Python using the Matplotlib and Seaborn libraries (Hunter, 2007; Waskom, 2021).

### 2.4 Machine Learning Models and Evaluation

The procedure for predicting the serogroup of a *Leptospira* isolate was designed as a two-step hierarchical classification pipeline. In the first step, each isolate was assigned to one of four major serological classes, hereafter referred to as seroclasses, as defined in previous work (Ferreira et al., 2024). In the second step, the isolate was classified into one of the serogroups belonging to the seroclass to which it had been assigned in the first stage. All classifiers were trained as binary Balanced Random Forest (BRF) models to determine whether a given isolate belonged to a specific seroclass or serogroup. In other words, each model was trained to discriminate between “member” and “non-member” instances for a given target group. The hyperparameters were set as follows: n_estimators=200, random_state=2, max_leaf_nodes=5, class_weight=“balanced”.

For the first-stage classification, models were trained using the filtered dataset, encompassing 384 selected genomes. For the second stage, separate BRF models were trained using only isolates corresponding to the serogroups within a single seroclass. Likewise, only the similarity data derived from proteins from representative samples of serogroups of that seroclass were included as input features for model training in this stage. Model evaluation was performed using distinct cross-validation schemes appropriate to the sample size of each classification level. The first-stage models were evaluated using 5-fold cross-validation, while the second-stage models were assessed using a leave-one-out (LOO) validation approach, chosen to account for the limited number of positive samples available for certain serogroups. The predictive performance of all models was evaluated based on accuracy, precision, recall, and F1-score metrics.

### 2.5 Reproducibility and data availability

All scripts, trained models, and feature matrices are available in a public GitHub repository (https://github.com/Spidey2004/Modelo_Leptospira). Genome accession numbers and associated serogroup metadata are listed in the Supplementary Tables S1 and S2.

## 3. Results

### 3.1 Dataset composition and exploratory analysis of the *rfb*-derived feature space

The feature matrix used to develop a machine learning model capable of serologically classifying pathogenic *Leptospira* isolates based on the gene composition of the *rfb* locus was comprised of 384 leptospiral samples (rows) and 549 proteins encoded in the *rfb* locus (columns). The selected samples were distributed across four major seroclasses and 22 distinct serogroups (Table 2). Among these, seroclass I was the most represented, encompassing 138 genomes assigned to nine serogroups. Within this seroclass, the serogroups *Icterohaemorrhagiae* (43 genomes) and *Pyrogenes* (28 genomes) were the most prevalent, followed by *Canicola* (23 genomes) and *Ballum* (19 genomes). Seroclass II comprised 129 genomes distributed across six serogroups, with *Pomona* (43 genomes) and *Grippotyphosa* (39 genomes) being the most represented. Seroclass III contained 86 genomes assigned to three serogroups, primarily from the *Sejroe* (39 genomes). Finally, Seroclass IV included 31 genomes, divided among three serogroups, with *Tarassovi* (18 genomes) as the most frequent representatives. Across all samples, *Leptospira interrogans* was the most represented species (199 genomes), followed by *L. borgpetersenii* (83 genomes), *L. kirschneri* (40 genomes), and *L. santarosai* (38 genomes). Among the 22 serogroups analyzed, 15 included isolates from at least two distinct species, indicating substantial interspecies distribution of serogroup classification. Only serogroups Ballum, Celledoni, Ranarum, Cynopteri, Djasiman, Panama, and Shermani were each represented by a single species. Similarly, among the nine *Leptospira* species included in the dataset, only *L. alstoni* and *L. mayottensis* were restricted to a single serogroup.

**Table 2:** Distribution of *Leptospira* isolates among classes, serogroups and species included in the machine learning analyses.

| Seroclass* | Serogroup | Lale | Lals | Lbor | Lint | Lkir | Lmay | Lnog | Lsan | Lwei | Total |
| --- | --- | --- | --- | --- | --- | --- | --- | --- | --- | --- | --- |
| I (138) | Ballum |  |  | 19 |  |  |  |  |  |  | 19 |
|  | Canicola |  |  |  | 22 | 1 |  |  |  |  | 23 |
|  | Celledoni |  |  |  |  |  |  |  |  | 1 | 1 |
|  | Icterohaemorrhagiae |  |  |  | 42 | 1 |  |  |  |  | 43 |
|  | Javanica | 1 |  | 13 |  |  |  |  | 1 |  | 15 |
|  | Manhao | 2 |  |  |  |  |  |  |  | 1 | 3 |
|  | Pyrogenes |  |  |  | 22 |  |  | 1 | 5 |  | 28 |
|  | Ranarum |  | 2 |  |  |  |  |  |  |  | 2 |
|  | Sarmin |  |  |  |  |  |  |  | 3 | 1 | 4 |
| II (129) | Australis |  |  |  | 14 |  |  | 3 |  |  | 17 |
|  | Autumnalis |  |  |  | 19 | 3 |  | 1 |  |  | 23 |
|  | Cynopteri |  |  |  |  | 2 |  |  |  |  | 2 |
|  | Djasiman |  |  |  | 2 |  |  |  |  |  | 2 |
|  | Grippotyphosa |  |  |  | 13 | 21 |  |  | 5 |  | 39 |
|  | Panama |  |  |  |  |  |  | 3 |  |  | 3 |
|  | Pomona |  |  | 3 | 30 | 10 |  |  |  |  | 43 |
| III (86) | Hebdomadis | 1 |  |  | 19 |  |  |  | 1 |  | 21 |
|  | Mini |  |  | 14 | 1 | 2 | 3 |  | 5 | 1 | 26 |
|  | Sejroe |  |  | 27 | 9 |  |  |  | 3 |  | 39 |
| IV (31) | Bataviae |  |  |  | 6 |  |  | 1 |  |  | 7 |
|  | Shermani |  |  |  |  |  |  |  | 6 |  | 6 |
|  | Tarassovi | 1 |  | 7 |  |  |  |  | 9 | 1 | 18 |
| Total |  | 5 | 2 | 83 | 199 | 40 | 3 | 9 | 38 | 5 | 384 |

\*total number of samples in each seroclass indicated in parentheses.
Lale: *L. alexanderi*; Lals: *L. alstonii*; Lbor: *L. borgpetersenii*; Lint: *L. interrogans*; Lkir: *L. kirschneri*; Lmay: *L. mayottensis*; Lnog: *L. noguchii*; Lsan: *L. santarosai*; Lwei: *L. weilii*.

A principal component analysis (PCA) was performed to investigate the overall distribution of *Leptospira* samples based on the gene composition of the *rfb* locus (Figure 1). The first two principal components explained the largest proportion of variance in the dataset and revealed clustering patterns largely consistent with the seroclass structure (Figure 1A). PCA was further applied separately to each seroclass to examine the distribution and clustering of their respective serogroups (Figure 1B–E). In most cases, serogroups formed well-defined and cohesive clusters. However, within seroclass II, serogroups such as Autumnalis, Pomona, and Grippotyphosa displayed greater dispersion and partial overlap, indicating reduced separation compared to other serogroups.

**Figure 1:**
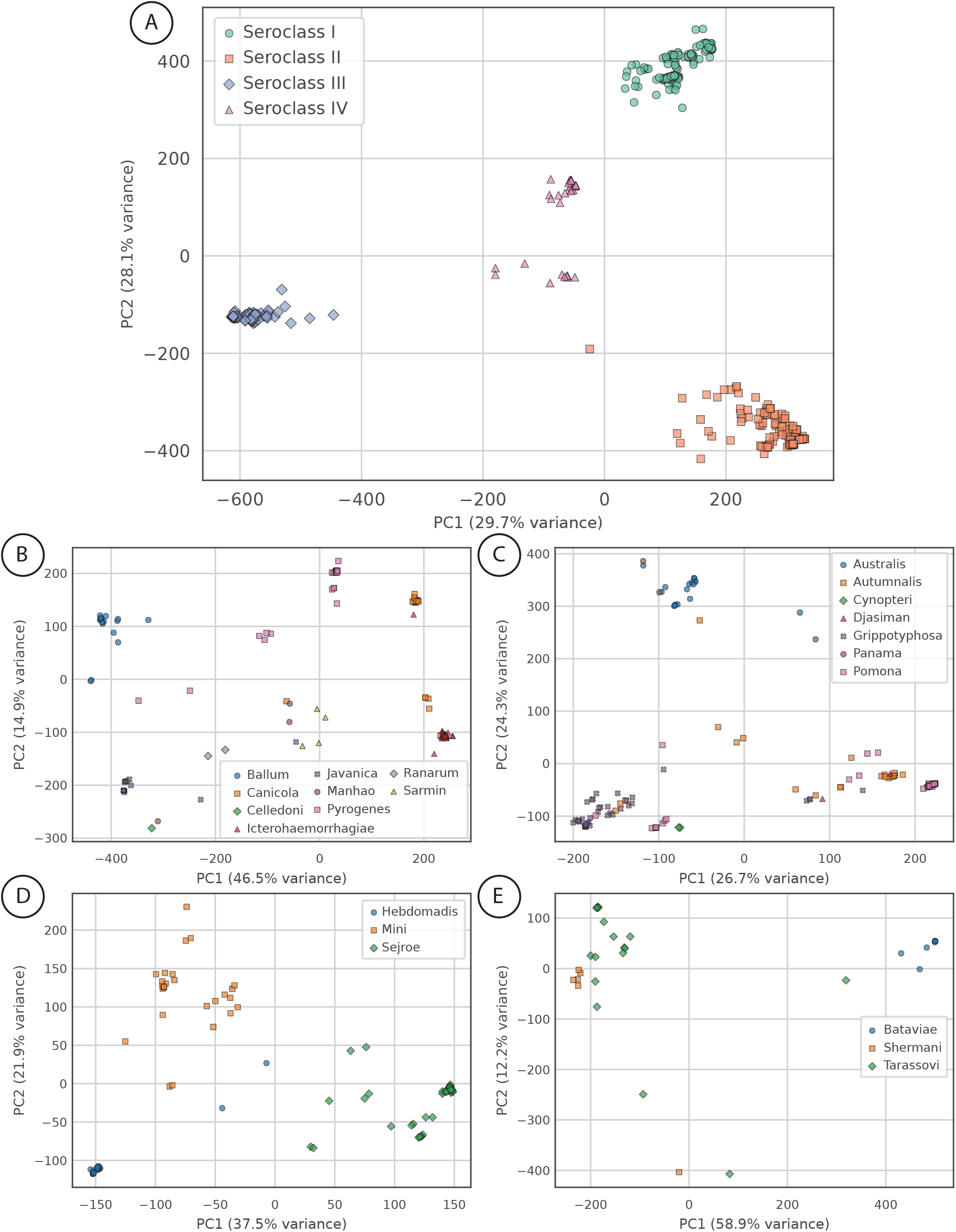
Principal Component Analysis (PCA) of *Leptospira rfb*-locus genomic features. (A) PCA including 384 *Leptospira* genomes representing four distinct seroclasses. (B–E) PCA performed independently for genomes assigned to seroclasses I (B), II (C), III (D), and IV (E). Each point represents a single genome. In panel A, colors and marker shapes denote seroclass membership, whereas in panels B–E they indicate serogroup classification within each seroclass.

### 3.2. *Leptospira* serogroup machine learning classifier

The procedure for predicting the serogroup of a *Leptospira* isolate was designed as a two-step hierarchical classification pipeline. In the first step, each isolate is assigned to one of four major seroclasses. Then, In the second step, the isolate is classified into one of the serogroups belonging to the seroclass to which it had been assigned in the first stage. Below, we present the analysis of the data and models generated in each stage.

#### 3.2.1. First stage classification (seroclass level)

In this stage, classifiers were generated to assign *Leptospira* isolates to one of the four seroclasses. For each seroclass, an independent binary classification model was trained using the complete dataset (384 samples), with targets indicating whether a sample belonged or not to that specific seroclass. This binary approach was chosen to account for potential cases in which a new isolate might not belong to any of the previously characterized seroclasses. A conventional multiclass model would force such samples into one of the existing seroclasses, potentially leading to misclassification.

After model training using the BRF algorithm, performance was evaluated through 5-fold cross-validation. The models correctly assigned all samples to their respective seroclasses, achieving perfect classification performance across all evaluated metrics. This result is consistent with the clear separation of seroclasses observed in the PCA analysis (Figure 1a), which indicated minimal inter-class overlap in the *rfb*-derived feature space.

#### 3.2.2. Second stage classification (serogroup level)

In the second stage of the classification pipeline, BRF models were trained to assign *Leptospira* isolates to specific serogroups within each of the four seroclasses. Each model was evaluated individually using the leave-one-out cross-validation strategy, and its performance was assessed through accuracy, precision, recall, and F1-score metrics (Table 3). Models for a few serogroups from seroclass I (Celledoni, Manhao, Ranarum, and Sarmin) and seroclass II (Cynopteri, Djasiman, and Panama) were not generated due to insufficient data for proper training.

**Table 3:**
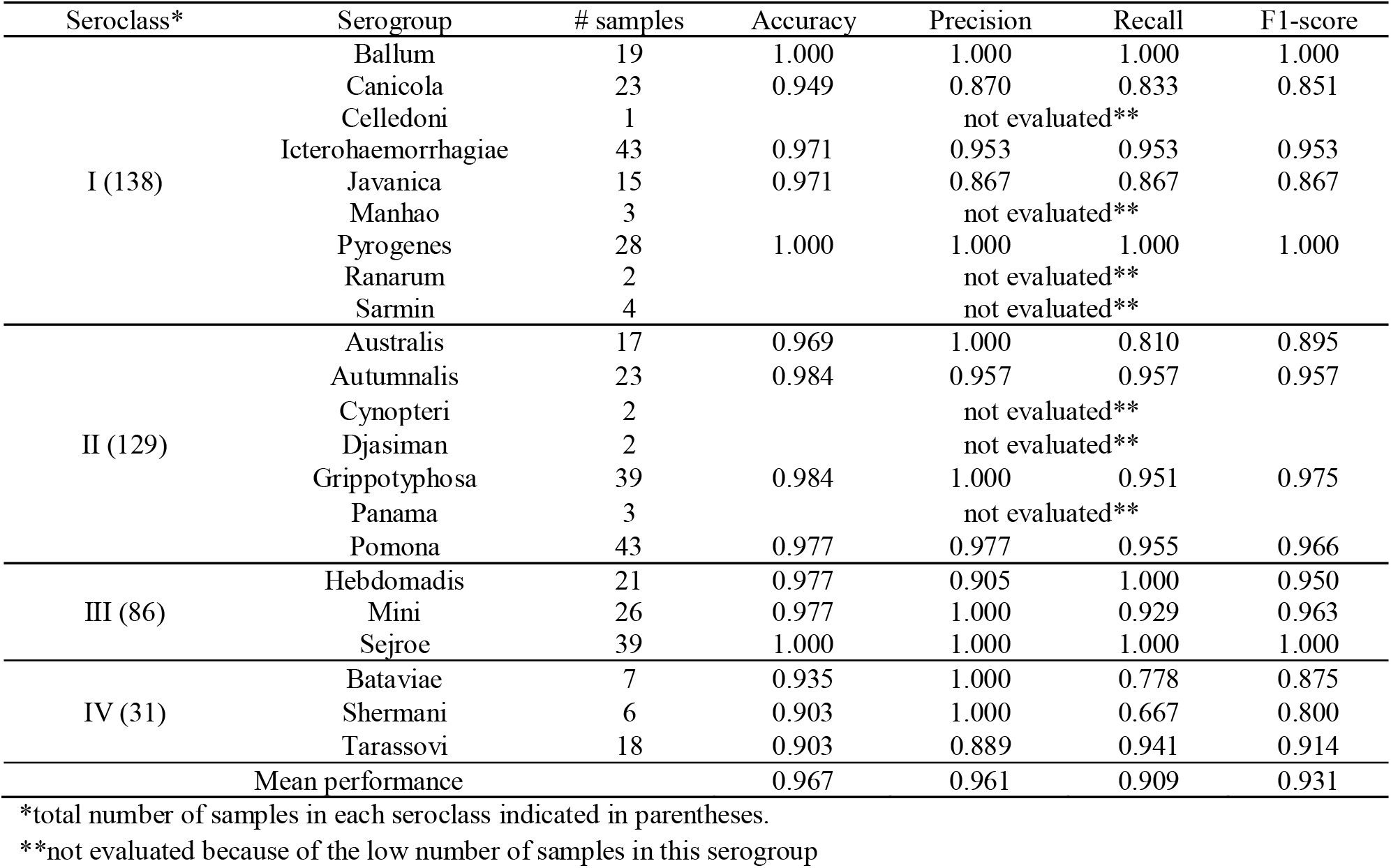
Performance of the models from the second-stage (serogroup classification)

Overall, the models achieved high predictive performance, with an average accuracy of 0.967, precision of 0.961, recall of 0.909, and F1-score of 0.931. Within seroclass I (n = 138), which contained the largest number of serogroups, the majority of models exhibited excellent performance, with F1-scores above 0.95 for serogroups Ballum, Icterohaemorrhagiae and Pyrogenes. Only the models for the Javanica and Canicola serogroup had a low F1-score compared to other models (above 0.85), but it was still considered high.

For seroclass II (n = 129), F1-scores ranged between 0.895 and 0.975, with the best results observed for the Grippotyphosa (F1 = 0.975), Autumnalis (F1 = 0.957) and Pomona (F1 = 0.966) serogroups. Slightly lower performance was recorded for the Australis serogroup, which yielded F1-scores of 0.895. seroclass III (n = 86) included serogroups Sejroe, Mini, and Hebdomadis, all of which achieved robust performance with F1-scores above 0.95, demonstrating consistent discriminative signal across these closely related serogroups. Finally, seroclass IV (n = 31), which contained fewer samples, showed more variable results, with F1-scores ranging from 0.8 for Shermani to 0.914 for Tarassovi.

A more detailed evaluation was performed by examining the prediction outcome and probability for each sample during leave-one-out cross-validation (Supplementary Table S5) across all models. A summary of these results is presented in Table 4, where rows represent the predictive models trained for individual serogroups, whereas columns correspond to the true serogroup assignment of the samples. The values indicate the number of samples predicted as positive in each category. Overall, the models predominantly assigned positive predictions to samples belonging to their respective serogroups, as evidenced by the higher counts observed along the main diagonal within each seroclass block. Off-diagonal entries represent cross-classifications, reflecting instances in which samples from one serogroup were predicted as positive by a model trained on another serogroup. These misclassifications were generally infrequent, supporting the overall discriminatory capacity of the models. However, notable exceptions were observed among serogroups represented by a limited number of samples. Specifically, all samples from the Celledoni, Cynopteri, and Panama serogroups were systematically misclassified as Javanica, Pomona, and Australis, respectively. This complete reassignment suggests a strong overlap in *rfb* locus gene composition between these pairs of serogroups, indicating reduced discriminatory signal within the selected genomic features. At the same time, the small sample size of these underrepresented serogroups likely contributed to this behavior by limiting the models’ ability to learn robust and distinctive patterns. Increasing the number of representative genomes for these serogroups would likely improve model calibration and enhance classification reliability.

**Table 4:** Distribution of samples classified as positive by each model across serogroups.

| Models |  | Number of positive samples |  |  |  |  |  |  |  |  |
| --- | --- | --- | --- | --- | --- | --- | --- | --- | --- | --- |
|  |  | Ball<br>(n=19) | Cani<br>(n=23) | Cell<br>(n=1) | Icte<br>(n=43) | Java<br>(n=15) | Manh<br>(n=3) | Pyro<br>(n=28) | Rana<br>(n=2) | Sarm<br>(n=4) |
| I | Ballum | 19 | 0 | 0 | 0 | 0 | 0 | 0 | 0 | 0 |
|  | Canicola | 0 | 20 | 0 | 4 | 0 | 0 | 0 | 0 | 0 |
|  | Icterohaemorrhagiae | 0 | 2 | 0 | 41 | 0 | 0 | 0 | 0 | 0 |
|  | Javanica | 0 | 0 | 1 | 0 | 13 | 1 | 0 | 0 | 0 |
|  | Pyrogenes | 0 | 0 | 0 | 0 | 0 | 0 | 28 | 0 | 0 |
| Models |  | Aust<br>(n=17) | Autu<br>(n=23) | Cyno<br>(n=2) | Djas<br>(n=2) | Grip<br>(n=39) | Pana<br>(n=3) | Pomo<br>(n=43) |  |  |
| II | Australis | 17 | 1 | 0 | 0 | 0 | 3 | 0 |  |  |
|  | Autumnalis | 0 | 22 | 0 | 1 | 0 | 0 | 0 |  |  |
|  | Grippotyphosa | 0 | 0 | 0 | 1 | 39 | 0 | 1 |  |  |
|  | Pomona | 0 | 0 | 2 | 0 | 0 | 0 | 42 |  |  |
| Models |  | Hebd<br>(n=21) | Mini<br>(n=26) | Sejr<br>(n=39) |  |  |  |  |  |  |
| III | Hebdomadis | 19 | 0 | 0 |  |  |  |  |  |  |
|  | Mini | 2 | 26 | 0 |  |  |  |  |  |  |
|  | Sejroe | 0 | 0 | 39 |  |  |  |  |  |  |
| Models |  | Bata<br>(n=7) | Sher<br>(n=6) | Tara<br>(n=18) |  |  |  |  |  |  |
| IV | Bataviae | 7 | 0 | 2 |  |  |  |  |  |  |
|  | Shermani | 0 | 6 | 3 |  |  |  |  |  |  |
|  | Tarassovi | 0 | 1 | 16 |  |  |  |  |  |  |

Ball: Ballum; Cani: Canicola; Cell: Celledoni; Icte: Icterohaemorrhagiae; Java: Javanica; Manh: Manhao; Pyro: Pyrogenes; Rana: Ranarum; Sarm: Sarmin; Aust: Australis; Autu: Autumnalis; Cyno: Cynopteri; Djas: Djasiman; Grip: Grippotyphosa; Pana: Panama; Pomo: Pomona; Hebd: Hebdomadis; Sejr: Sejroe; Bata: Bataviae; Sher: Shermani; Tara: Tarassovi.

### 3.3. Relevant genes for serogroup classification

Because we employed a tree-based algorithm (Random Forest), it was possible to identify which genes (features) were most frequently used in the decision-making process across the ensemble of trees composing each model. In this analysis, proteins exhibiting feature importance above 0.025 were considered to be relevant features for the model. This threshold retained approximately 2% of the features in the first-stage models and 3–12% in the second-stage models. In total, 52 and 114 distinct proteins were identified as important for classification in the first- and second-stage models, respectively (Supplementary Table S6).

For the serogroup classification, by projecting the genomic positions of the most informative genes onto the *rfb* locus of representative strains from each serogroup (Figure 2), we observed that the majority of high-importance features clustered within the first half of the locus. This non-random distribution suggests that the proximal region of the *rfb* locus concentrates key genetic determinants underlying serogroup specificity. The functional composition of the proteins identified as important features reveals that most of these proteins are associated with carbohydrate biosynthesis and modification pathways. These include glycosyltransferases, class I SAM-dependent methyltransferases, aminotransferases of the DegT/DnrJ/EryC1/StrS family, and multiple oxidoreductases such as SDR family proteins and NAD-dependent epimerase/dehydratases, which collectively contribute to the structural diversity of surface-exposed polysaccharides. For some serogroups, including Sejroe, Pomona, Bataviae, and Shermani, discrimination was driven by highly specific markers that were consistently present in isolates belonging to these serogroups but largely absent in others. In contrast, for serogroups such as Pyrogenes, Australis and Grippotyphosa, classification performance was primarily supported by the consistent absence of particular genes in the ingroup coupled with their high prevalence among other serogroups. Notably, some proteins emerged as important features across multiple serogroup-specific models. These observations indicate that discrimination is not solely dependent on uniquely serogroup-specific genes. Rather, they support a model in which serogroup differentiation is driven by combinatorial patterns of gene presence and absence within the *rfb* locus.

**Figure 2.**
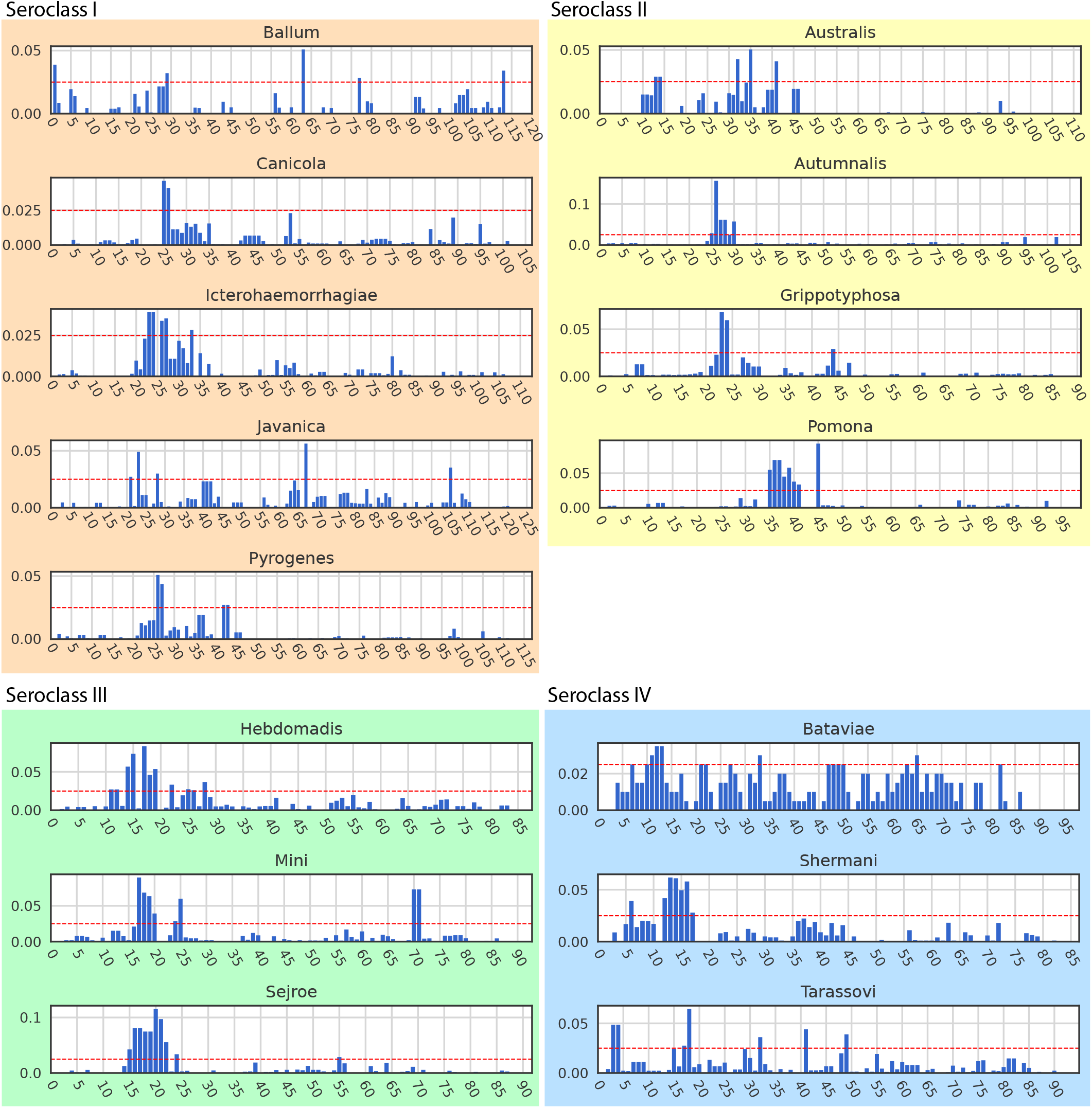
Feature importance profiles for each serogroup-specific model. Each panel represents one serogroup model. The y-axis indicates the feature importance values assigned by the model, whereas the x-axis corresponds to the genomic position of genes along the *rfb* locus. Higher importance values indicate genes that contributed more strongly to the classification decision. The red horizontal line in all panels represents the importance threshold of 0.025 adopted for highlighting relevant features. Only features associated with gene presence are displayed; genes for which discriminatory power was driven by their characteristic absence in a given serogroup are described in Supplementary Table S6.

### 3.4. Final serogroup classification framework and its validation

Based on the subset of proteins identified as relevant for classification, new machine learning models were trained using only these selected features. When evaluating their performance using the same strategies described above, the reduced models achieved performance comparable to that of the full-feature models across all metrics, with perfect score for the models of first stage and a mean F1-score of 0.948 for the models of second stage. These results indicate that a reduced set of informative genes is sufficient to retain the predictive signal required for accurate classification. Therefore, the reduced models were selected to constitute the final classification framework.

To assess generalization performance, an independent validation dataset comprising 30 genome samples not used during model training was analyzed. The prediction results are summarized in Table 5. Overall, the framework demonstrated high predictive accuracy, with most genomes correctly assigned to their respective serogroups and consistently high prediction probabilities, frequently exceeding 0.90. Correct classifications were observed across multiple pathogenic species, including *L. interrogans, L. kirschneri*, and *L. borgpetersenii*, indicating that the genomic signatures captured from the *rfb* locus are robust and transferable across distinct evolutionary lineages.

**Table 5:** Classification of leptospiral samples from validation dataset.

| Acession | Species | Strain | True serogroup | Prediction | Prediction probability |
| --- | --- | --- | --- | --- | --- |
| GCF_041647705.1 | *L. kirschneri* | H9 | Australis | Australis | 78.50% |
| GCF_041647755.1 | *L. interrogans* | H44 | Pomona | Pomona | 100.00% |
| GCF_045159415.1 | *L. interrogans* | RJ19115 | Icterohaemorrhagiae | Icterohaemorrhagiae | 96.60% |
| GCF_046529585.1 | *L. interrogans* | Ballico | Australis | Australis | 98.50% |
| GCF_046848885.1 | *L. interrogans* | Jez Bratislava | Australis | Australis | 99.00% |
| GCF_046848895.1 | *L. interrogans* | Van Tienen | Bataviae | Bataviae | 98.00% |
| GCF_046848905.1 | *L. interrogans* | Djasiman | Djasiman | Grippotyphosa | 68.50% |
| GCF_048973995.1 | *L. borgpetersenii* | BU-20 | Sejroe/Hardjo | Sejroe | 100.00% |
| GCF_048974015.1 | *L. borgpetersenii* | BU-18 | Sejroe | Sejroe | 100.00% |
| GCF_048974035.1 | *L. borgpetersenii* | BU-19 | Sejroe | Sejroe | 100.00% |
| GCF_048974055.1 | *L. borgpetersenii* | BS-39 | Sejroe | Sejroe | 100.00% |
| GCF_049953905.1 | *L. interrogans* | USDA MIL-020 | Mini | Mini | 100.00% |
| GCF_049954495.1 | *L. interrogans* | USDA HBL-010 | Hebdomadis | Hebdomadis | 100.00% |
| GCF_049954505.1 | *L. borgpetersenii* | USDA SJL-010 | Sejroe | Sejroe | 100.00% |
| GCF_049955055.1 | *L. interrogans* | USDA ARL-050 | Australis | Australis | 99.00% |
| GCF_049955065.1 | *L. borgpetersenii* | USDA TAL-010 | Tarassovi | Tarassovi | 100.00% |
| GCF_049955695.1 | *L. interrogans* | USDA PYL-010 | Pyrogenes | Pyrogenes | 100.00% |
| GCF_049955705.1 | *L. interrogans* | USDA POL-010 | Pomona | Pomona | 100.00% |
| GCF_049956375.1 | *L. interrogans* | USDA ICL-020 | Icterohaemorrhagiae | Icterohaemorrhagiae | 96.60% |
| GCF_049956385.1 | *L. interrogans* | USDA SJL-060 | Sejroe | Sejroe | 100.00% |
| GCF_049956975.1 | *L. interrogans* | USDA GRL-020 | Grippotyphosa | Grippotyphosa | 95.80% |
| GCF_049957005.1 | *L. interrogans* | USDA CAL-010 | Canicola | Canicola | 99.90% |
| GCF_049957935.1 | *L. interrogans* | USDA BTL-020 | Bataviae | Bataviae | 98.00% |
| GCF_049957945.1 | *L. borgpetersenii* | USDA BML-011 | Ballum | Ballum | 100.00% |
| GCF_049958755.1 | *L. interrogans* | USDA ATL-010 | Autumnalis | Autumnalis | 100.00% |
| GCF_049958765.1 | *L. interrogans* | USDA ARL-010 | Australis | Australis | 98.50% |
| GCF_052161685.1 | *L. interrogans* | L7 | Icterohaemorrhagiae | Icterohaemorrhagiae | 99.50% |
| GCF_052161695.1 | *L. interrogans* | L4 | Canicola | Canicola | 99.90% |
| GCF_052161705.1 | *L. interrogans* | L2 | Canicola | Canicola | 99.90% |
| GCF_052161715.1 | *L. interrogans* | L3 | Canicola | Canicola | 99.90% |

A single discrepancy was observed for strain Djasiman (serogroup Djasiman), which was predicted as Grippotyphosa. This misclassification is attributable to the limited representation of the Djasiman serogroup in the training dataset (n = 2), which prevented the development of a dedicated classification model for this serogroup and restricted the learning of distinctive genomic patterns. Consequently, Djasiman isolates were classified based on their similarity to models trained for other serogroups. Notably, the relatively low prediction probability assigned to this sample (68.5%) indicates classification uncertainty and may serve as a useful indicator for flagging potential misclassifications in practical applications.

## 4. Discussion

In this study, we proposed and validated a genomic-based machine learning framework for serogroup classification in *Leptospira*. To the best of our knowledge, this is the first study to apply machine learning methods to develop predictive classifiers specifically targeting serogroup-level identification in this genus based solely on genomic data. By leveraging the antigenic diversity encoded in the *rfb* locus, which underlies genes involved in the biosynthesis of the O-antigen component of lipopolysaccharides (LPS) (Mitchison et al., 1997), our models effectively captured the genetic determinants that distinguish major serological groups. The two-step hierarchical strategy, first classifying samples into four broader antigenic classes and subsequently refining the prediction at the serogroup level, demonstrated robust performance, with perfect score for the first stage and average F1-scores of 0.948 for second stage. These values indicate that the genomic features derived from the *rfb* locus encode sufficient discriminatory power to accurately predict serological identity at serogroup level. In this sense, our results provide genomic validation of the long-assumed link between *rfb* organization and serological phenotype (Adler and de la Peña Moctezuma, 2010; Bulach et al., 2000; de la Peña-Moctezuma et al., 1999; Lehmann et al., 2014), bridging classical and molecular serological classification.

An important consideration when interpreting the misclassification for some models concerns the accuracy of serological annotations used for model training. Because traditional methods such as MAT and CAAT are susceptible to cross-reactivity and subjective interpretation (Adler and de la Peña Moctezuma, 2010; Levett, 2001), inconsistencies in serogroup labeling may be present in public databases. Such misclassifications introduce noise into the training dataset, which can artificially limit the model’s performance or produce apparent misclassifications. Therefore, the performance metrics reported here likely represent a lower bound of the true predictive potential of genomic data, emphasizing the importance of improving and standardizing serological metadata for future studies.

Despite its promising results, this study has some limitations. First, the representation of certain serogroups remains limited in public genomic databases, which may restrict model generalizability, particularly for rare or newly emerging variants. Second, although the binary classification strategy reduces the risk of forcing isolates into predefined categories, it underscores the need for complementary mechanisms capable of detecting potentially novel serogroups that fall outside established classifications. Future research may address these limitations through the continued expansion of whole-genome sequencing efforts for *Leptospira* isolates, conducted in parallel with standardized serological characterization, and the incorporation of unsupervised clustering approaches, which could facilitate the detection of previously unrecognized antigenic patterns and contribute to the refinement of the proposed hierarchical classification structure.

Since the proposed models rely on genomic sequence data as input, such information may not be readily available in routine clinical settings, potentially limiting their immediate applicability in diagnostic workflows. However, the analysis of feature importance provides an important translational advantage. In particular, genes consistently exhibiting high importance and strong serogroup specificity could serve as candidates for the development of PCR-based diagnostic tools, enabling rapid and cost-effective serological classification without the need for whole-genome sequencing. Such an approach could complement and refine previously proposed molecular typing strategies (Cai et al., 2010; Wenderlein et al., 2024) by providing a data-driven framework for marker selection grounded in machine learning–based feature prioritization. Therefore, beyond predictive performance, the feature importance analysis offers a pathway toward improving and modernizing molecular diagnostic methods for *Leptospira* serogroup classification.

Finally, we propose the term “seroclass” to designate a higher-order grouping of *Leptospira* serogroups. Serogroups within the same seroclass exhibit greater gene composition similarity at the *rfb* locus and frequent antigenic cross-reactions than those in different groups, indicating an intermediate hierarchical level not captured by the current serological framework. This terminology refines previous use of broader terms such as “cluster” (Chinchilla et al., 2023) or “class” (Ferreira et al., 2024) and provides a clearer structure for interpreting serological diversity.

## 5. Conclusion

In conclusion, our results demonstrate that machine learning models trained on *rfb*-derived genomic features can accurately reproduce the traditional serological classification of *Leptospira*. This approach offers a scalable, standardized, and reproducible alternative to serological assays, and contributes in paving the way toward a genomics-informed system for classification and surveillance of this complex and diverse bacterial genus.

## Acknowledgements

We would like to thank the teams from Bioinformatics Multidisciplinary Environment (BioME/IMD) at UFRN, Centro de Processamento de Alto Desempenho (CEPAD/ICB) at UFMG, and High-Performance Computing Center (NPAD) at UFRN for the support on the computational resource. We are also grateful to NCBI and the Institut Pasteur teams for the curation and maintenance of the data used in this work.

## Data statement

All genomic data used in this work are hosted in NCBI and BIGSdb, which are public databases.

## Declaration of competing interest

The authors declare that they have no competing interests.

## Supplementary Tables

**Supplementary Table S1**: Metadata for the 721 *Leptospira* strains included in this study. **Supplementary Table S2**: Metadata for the *Leptospira* strains included in the validation dataset. **Supplementary Table S3**: Gene identity feature matrix derived from the *rfb* locus.

**Supplementary Table S4**: Pairwise correlation matrix among *Leptospira* strains based on *rfb* locus gene profiles.

**Supplementary Table S5**: Serogroup classification probabilities for each sample during leave-one-out cross-validation.

**Supplementary Table 6**: High-importance features for each seroclass- and serogroup-specific model

## Notes

https://github.com/Spidey2004/Modelo_Leptospira

### Competing Interest Statement

The authors have declared no competing interest.

### Summary of Updates

Final machine learning models updated accounting only the relevant genes.

